# BCR analysis of single-sorted, putative IgE^+^ memory B cells in food allergy: *an ephemeral existence?*

**DOI:** 10.1101/510743

**Authors:** Rodrigo Jiménez-Saiz, Yosef Ellenbogen, Kelly Bruton, Paul Spill, Doron D. Sommer, Hermenio Lima, Susan Waserman, Sarita U. Patil, Wayne G. Shreffler, Manel Jordana

**Author notes:** Equal contributions (Rodrigo Jiménez-Saiz, Yosef Ellenbogen, Kelly Bruton). Correspondence to: Manel Jordana, MD, PhD. Dept. of Pathology & Molecular Medicine, McMaster Immunology Research Centre, McMaster University, MDCL 4013, 1280 Main St W, Hamilton, ON, L8S 4K1, Canada. Vaccines & Dendritic Cells Lab, Department of Biochemistry and Molecular Biology, Chemistry School, Complutense University, 28040 Madrid, Spain (Rodrigo Jiménez-Saiz).

## Abstract

Immunoglobulin (Ig) E is the critical effector molecule in allergic reactions. Consequently, research efforts to understand the biology of IgE-expressing cells is of paramount importance. In particular, the role of IgE^+^ memory B cells (MBCs) in the perpetuation of allergic reactivity has been the subject of intense research. Studies in mice have convincingly established that IgE^+^ B cells are rare and transient and, therefore, an unlikely candidate to maintain allergic disease. In contrast, IgE^+^ MBCs have been detected by flow cytometry in the sputum and peripheral blood of humans and have been proposed as a clinical marker of allergic disease. We established a method to genetically validate, at the single-cell level, the putative IgE^+^ MBCs identified by flow cytometry from humans. We, then used this information to develop an enhanced flow cytometry protocol that more accurately identifies *bona fide* IgE^+^ MBCs. We found that IgE^+^ MBCs were detected in some patients with atopic dermatitis, but at a frequency that was ~100 times lower than previously reported. We also found that IgE^+^ MBCs were undetectable in PBMCs from peanut allergic patients. These findings provide tools to identify *bona fide* IgE^+^ MBCs, demonstrate their extreme rarity in circulation and are consistent with the lack of a central role for IgE^+^ MBCs in the maintenance of allergic sensitivity.

**One Sentence Summary:** The frequency of IgE^+^ MBCs in the peripheral circulation of humans is orders of magnitude lower than previously reported and comparable between allergic and healthy donors, which cautions about the clinical utility of their assessment.

## Introduction

About 250 million people have food allergies worldwide and suffer clinical manifestations that range from mild local symptoms to life-threatening systemic anaphylaxis (*1*). In the USA alone, there are over 30,000 food-induced anaphylactic episodes each year (*2*), and ~200,000 emergency department visits for food-related acute reactions (*3*). Of particularly significant concern are allergies to peanuts (PN), tree nuts, fish and shellfish because they are lifelong in the majority of patients. This situation is aggravated by the lack of curative treatments. In fact, allergen-avoidance, together with the administration of epinephrine upon the onset of a systemic reaction, is the standard of care (*4*). The critical effector molecule in food-induced allergic reactions is immunoglobulin (Ig) E (*5*). Consequently, research efforts to understand the biology of IgE and the mechanisms of IgE memory are of paramount importance, and central to that is the accurate identification of IgE-expressing B cells (*6–8*).

The identification of IgE^+^ B cells *in vivo* is hindered by other leukocytes that can bind IgE through high- (FcεRI) or low- (FcεRII/CD23) affinity receptors (*9, 10*). The frequency of populations such as mast cells, basophils, dendritic cells, as well as B cells, that stain positive for IgE though they do not express it, is usually several times higher than IgE-expressing B cells (*11*). In addition, the density of IgE-BCRs is relatively low as compared to other BCRs, making the detection of IgE-expressing B cells exceedingly difficult (*12*). To identify *bona fide* IgE-expressing B cells, several staining methods have been reported. These include the removal of receptor-bound IgE *via* acid washes (*13*), the step-wise exclusion of B cells that stain for BCRs other than IgE (*14*), and the blockade of surface IgE to exclusively detect intracellular IgE (*15*).

In addition to technical challenges, the biology of IgE^+^ B cells remains enigmatic (*8, 16*). Studies in mouse models of Th2 immunity against helminths, haptens, and food allergens have reported populations of IgE-expressing germinal center B cells and plasma cells (*15, 17–20*), while IgE^+^ memory B cells (MBCs) were undetected. Furthermore, IgE^+^ B cells are predisposed to differentiate into short-lived plasma cells (*15*), and autonomous IgE-BCR signalling promotes apoptosis (*21*), thus limiting their survival. The body of data from murine models demonstrates that IgE^+^ MBCs are extremely rare, at best, therefore suggesting that IgE-mediated recall responses are derived from non-IgE expressing MBCs, particularly IgG1(*8, 22, 23*). In stark contrast, several human studies have claimed that a population of IgE^+^ MBCs can be identified in peripheral blood mononuclear cells (PBMCs) of healthy, atopic and food-allergic donors (*14, 24–26*), as well as in the sputum of asthmatics (*27, 28*). Moreover, the presence of these cells in circulation has been proposed as a prognostic marker of allergy (*24*). It is unclear whether the discrepancy between mouse and human data is due to fundamental differences between species in IgE^+^ B cell biology or to caveats in the method used for the identification of IgE^+^ MBCs. Therefore, a critical analysis and validation of the methods used to identify these cells in humans is of the utmost biological and clinical relevance.

Herein we describe a method to genetically validate, at the single-cell level, the flow cytometric identification of putative human IgE^+^ MBCs. BCR amplification of single cells isolated from peanut allergic patients that were identified as IgE-expressing cells using previously reported staining methods, as well as non-allergic donors, demonstrated a non-IgE identity in all instances. Using *in vitro* class switched cells, we then established a novel flow cytometry method to identify *bona fide* IgE^+^ MBCs. Single cell molecular validation was carried out in human PBMCs. IgE^+^ MBCs were detected only in PBMCs from 2 of 4 patients with atopic dermatitis, at a frequency 0.0015% from total B cells, which is ~100 times lower than previously reported (*14*). No IgE^+^ MBCs were detected in PBMCs from 9 PN-allergic patients and 10 non-allergic controls using our enhanced, genetically-validated, flow cytometry method. This research provides a genetically validated tool to identify *bona fide* IgE^+^ MBCs, demonstrates their extreme rarity, and questions the clinical utility of their detection in the peripheral circulation.

## Results

### Method to single-sort IgE^+^ MBCs and genetically validate their identity

The divergent findings on the existence of IgE-expressing MBCs between mice and humans could be partly due to the precision of techniques used in their quantification. While the identification of IgE-expressing MBCs cells in mice have been comprehensive, the identification of these cells in humans has relied on flow cytometry. In light of this, we sought to establish a method to validate the flow cytometric identification of IgE^+^ MBCs via genetic analysis of the BCR isotypes from single-sorted cells. We modified a single-cell B cell PCR amplification method (*29*) to amplify the variable region of IgE B cell heavy chain transcripts with primers shown specific to the 5’ end of IgE heavy chain constant region (*14*). This technique requires a single cell-nested RT-PCR and subsequent DNA gel electrophoresis for visualization of amplicons (**Fig. 1A**). Then amplificons undergo Sanger sequencing and resulting sequences are aligned to human heavy chain constant regions alleles of all antibody isotypes to assess homology. To evaluate that the PCR technique can amplify IgE transcripts with high specificity and sensitivity, we single-sorted a human IgE-expressing cell line (U266; ATCC TIB-196) and performed IgE RT-PCR following the same methodology (**Fig. 1B**).

**Fig. 1.**
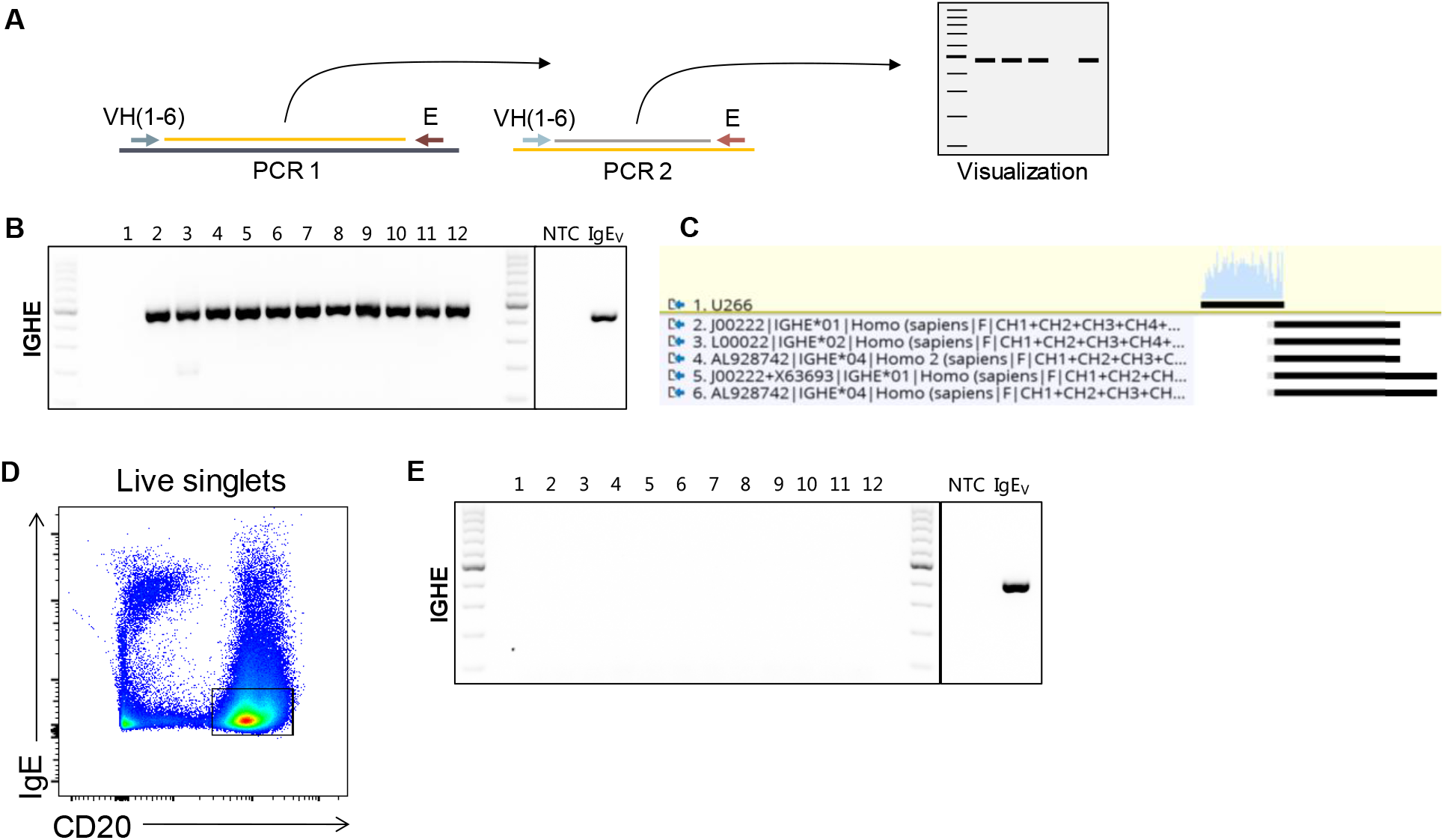
Methodology to validate genetically the IgE identity of single-sorted cells. **(A)** Schematic of the single-cell B cell PCR amplification method to amplify the variable region of IgE B cell heavy chain transcripts with primers shown specific to the 5’ end of IgE heavy chain and subsequent DNA gel electrophoresis. **(B)** Single-sorted human IgE-expressing myeloma cells were assayed by nested PCR; **(C)** the IgE transcripts were Sanger-sequenced and shown to align with IgE constant region (IGHE) alleles. **(D)** Peripheral blood CD20^+^ B cells that stained negative for IgE were single sorted and **(E)** IgE transcripts did not amplify as evidenced by DNA gel electrophoresis.

Our protocol demonstrated high sensitivity, amplifying on average over 90% of cells tested, the sequences of which all aligned to IgE constant region (IGHE) alleles (**Fig. 1C**). Additionally, we generated a DNA vector containing a human IgE backbone (IgEV) for use as a positive control. To determine that our technique specifically amplifies IgE transcripts and no other isotypes, we single-sorted peripheral blood B cells (CD20^+^) that did not stain for IgE (**Fig. 1D**). Using the same RT-PCR strategy as in **Fig. 1A**, none of the IgE^−^ B cells amplified (**Fig. 1E**), indicating that the technique is both sensitive and specific. Together, these data demonstrate that this system accurately amplifies rearranged IgE heavy chain variable sequences, specifically in single-sorted IgE-expressing cells.

### Detection of spurious IgE^+^ MBCs by current flow cytometric methods

Reported frequencies of IgE-expressing MBCs in peripheral blood vary depending on the flow cytometric identification strategies (*14, 24, 27, 28, 30, 31*). A basic approach to identify these cells involves intracellular staining of IgE without preventing staining of cytotropic IgE (*27, 28*). A more stringent detection method was reported by Berkowska *et al*.(*14*), which involves the stepwise exclusion of each BCR isotype via extracellular staining. Hereinafter, we refer to these IgE detection methods as basic IgE staining and step-wise exclusion, respectively. With our single-cell IgE amplification protocol, we sought to validate the frequency of peripheral blood IgE-expressing MBCs using these previously reported flow cytometric approaches. Briefly, PBMCs were isolated from PN-allergic donors. As B cells comprise ~3-15% of PBMCs in adults (*32*) we reasoned that B cell enrichment would empower the analysis of the MBC compartment. Therefore, B cells from PBMCs were purified (purity ~80%) via negative selection and stained for flow cytometry using the basic IgE staining and the step-wise exclusion methods.

MBCs were identified as live singlets CD20^+^CD38^lo-med^CD27^+^ and IgD^−^IgM^−^ and IgE^+^ MBCs were further identified through basic IgE staining or the step-wise exclusion approach (**Fig. 2B**). Twelve putative IgE-expressing MBCs were single-cell sorted from each staining technique for subsequent single-cell nested PCR. Basic IgE staining revealed a 3.4% frequency of IgE^+^ MBCs cells from B cells. However, no cells amplified with IgE-specific RT-PCR (**Fig. 2C**). The step-wise exclusion approach reported a frequency of putative IgE-expressing MBCs 20 times lower, 0.17% but, likewise, none of the cells amplified with IgE-specific RT-PCR (**Fig. 2D**). To delineate the BCR identity of the spurious cells that fell into the IgE gate we generated a cocktail of primers specific to IgG, IgA, and IgM heavy chain regions (GAM). Using our single-cell nested PCR strategy, over 90% of the sorted cells amplified with GAM primers and the rest did not amplify (<10%), which is consistent with previous reported efficiencies of these primers (*33*). Amplicons were confirmed to align predominantly with IGHG and to a minor extent to IGHA, or IGHM through Sanger sequencing. These data demonstrate that previously reported IgE^+^ MBC flow cytometry detection protocols result in a high rate of false-positive events.

**Fig. 2.**
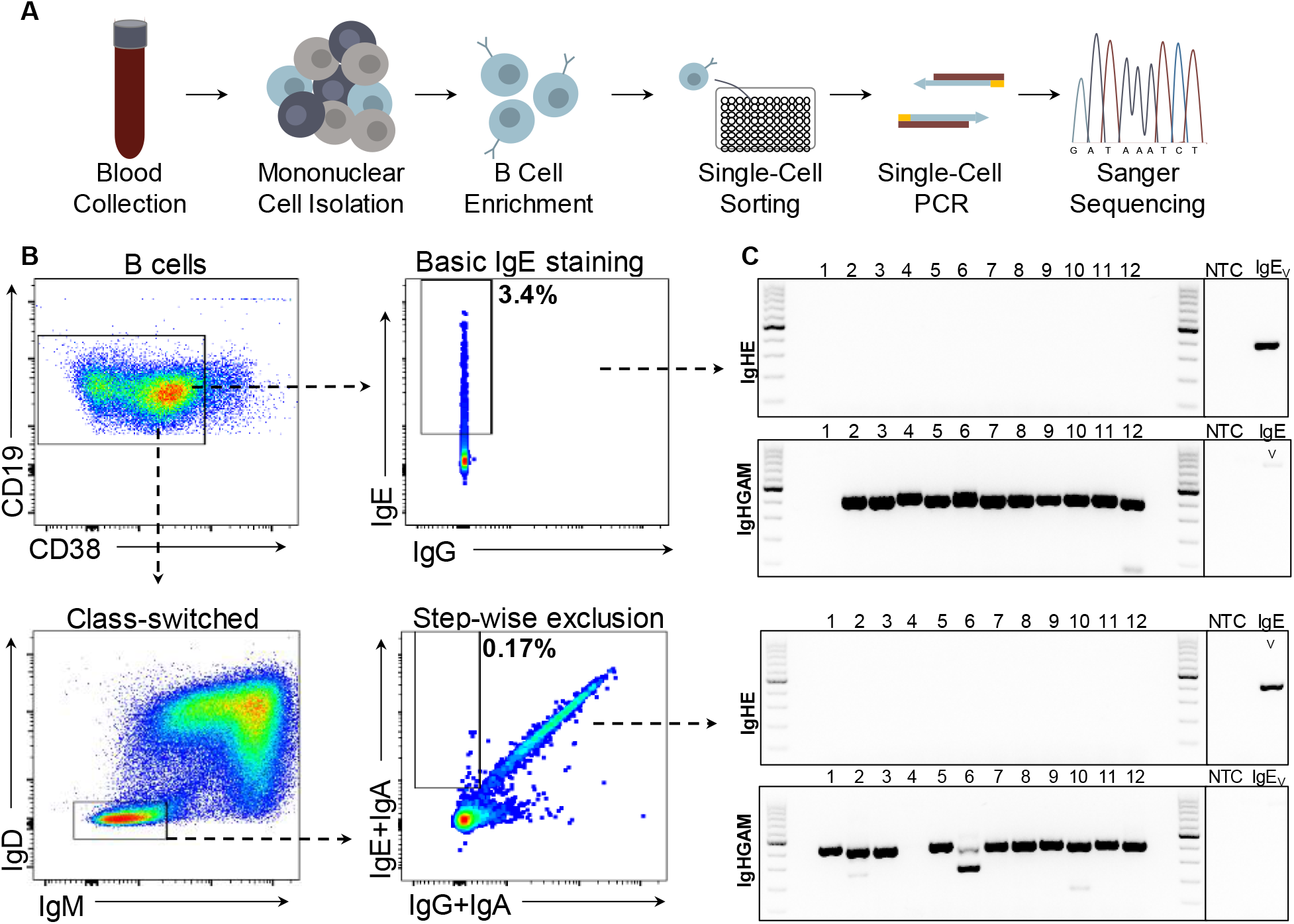
BCR analysis of single-sorted putative IgE^+^ MBCs stained with commonly used methods demonstrates a non-IgE identity. **(A)** Schematic of the experimental design to purify B cells from human blood, single-sort putative IgE^+^ MBCs following different staining methods and genetic validation of their BCR identity. **(B)** Gating strategy employed to single-sort putative IgE^+^ MBCs stained with a basic IgE staining (upper) or step-wise exclusion (lower) method. **(C)** BCR amplification with primers specific for IgE (IgHE) or a mix specific of IgG, IgA and IgM (IgHGAM) of single-sorted cells stained with different methods. Data are representative of 5 independent experiments (1-2 donors per experiment and 12-24 cells single-sorted per donor).

### Development of enhanced flow cytometric method for detection of IgE^+^ MBCs

The demonstration that previously reported flow cytometric methods for IgE^+^ MBC identification are faulty due to IgM^+^, IgA^+^ and IgG^+^ MBC contamination in the putative IgE gate, prompted us to ascertain the true frequency of these cells. Since the step-wise exclusion method generated a substantially lower number of spurious events than the basic IgE staining method, we sought to modify the former to remove contamination from non-IgE-expressing cells, which largely originated from IgG^+^ memory B cells carrying cytotropic IgE (data not shown). This enhanced protocol consisted of purifying B cells from PBMCs and a step-wise exclusion of IgD^+^, IgM^+^, IgA^+^, IgG^+^ (**Fig 3A**). Furthermore, the use of a polyclonal anti-IgG antibody markedly contributed to resolve the double negative population of MBCs (IgD^−^IgM^−^IgG^−^IgA^−^IgE^−^) compared to the previous step-wise exclusion method (3.8% *vs*. 58.6%). As B cells canonically express at least one BCR isotype, we reasoned that the population arose from B cells with a low BCR surface density, in which the use of a polyclonal (rather than monoclonal) antibody to stain for surface IgG was superior. Notably, the improved staining resulted in a significantly lower frequency of cells in the IgE gate, at 0.006% of total B cells, which was considered background as it was comparable to the frequency observed in the FMO (0.01% of total B cells; **Fig. 3A**).

**Fig. 3.**
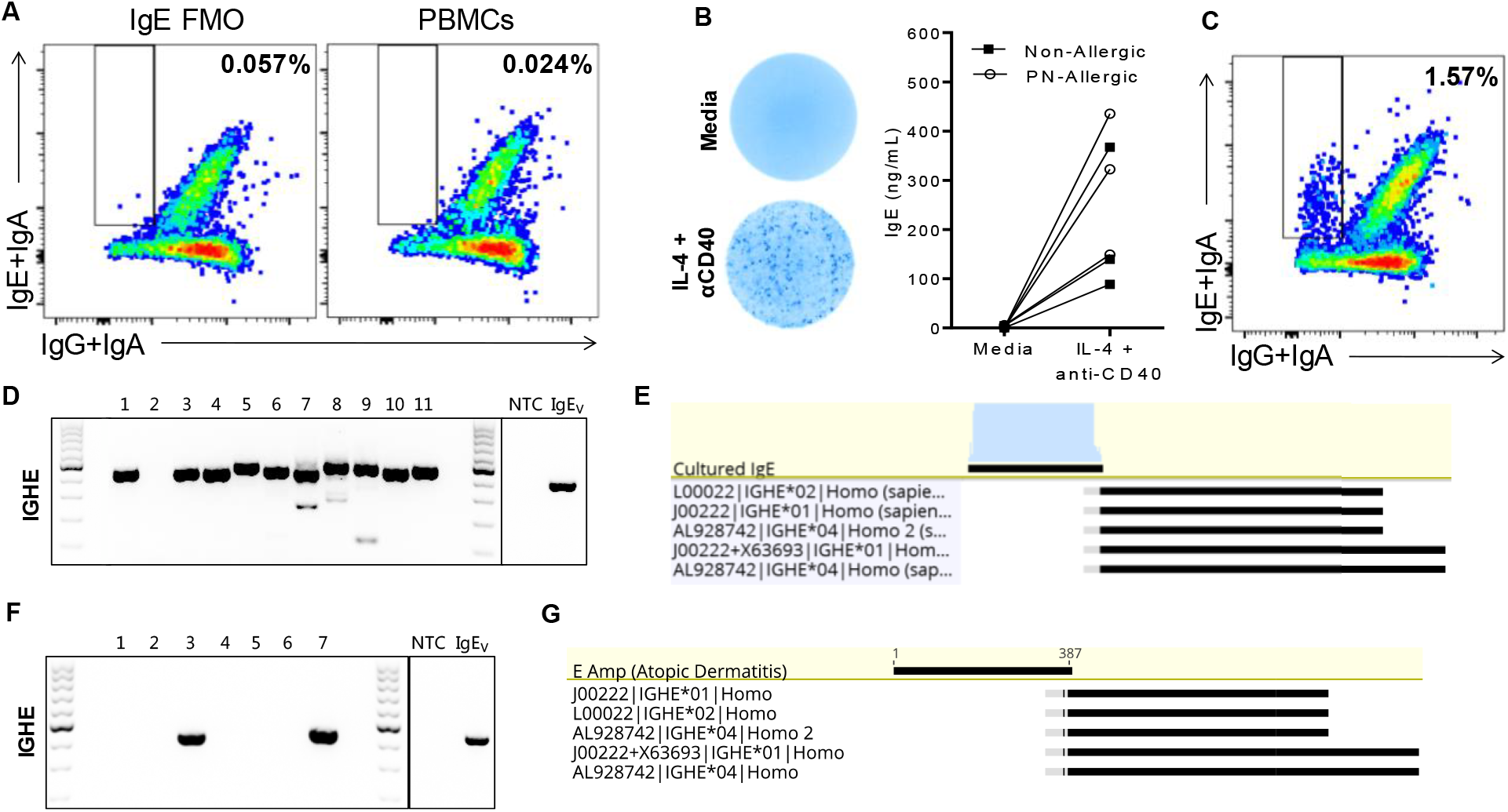
**(A)** Development of an enhanced flow cytometric method for detection of IgE^+^ MBCs. **(A)** Cytometric detection of IgE^+^ MBCs in healthy donors with the enhanced method. **(B)** PBMCs cultured with media or IL-4 and anti-CD40 were analyzed by ELISPOT and ELISA to detect IgE-secreting cells and secreted IgE respectively. **(C)** IgE^+^ MBCs in PBMCs stimulated with IL-4 and anti-CD40 were single-sorted; **(D)** amplification of IgE transcripts was assessed in agarose gels and **(E)** as well as their alignment to the constant region of IgHE following Sanger sequencing. Data are representative of 2 independent experiments (1-2 donors per experiment and 12-24 cells single-sorted per donor).

To validate that our enhanced staining technique was capable of detecting IgE^+^ MBCs, we stimulated PBMCs in culture with IL-4 + anti-CD40, which is known to induce IgE class-switching (*34*). As expected, culturing under these conditions resulted in the robust emergence of IgE-secreting cells and IgE as detected by total IgE ELISPOT and ELISA, respectively (**Fig. 3B**). A population of putative IgE^+^ MBCs was observed through the enhanced step-wise exclusion method (**Fig. 3C**) and their IgE-identity was confirmed through single-cell nested RT-PCR and Sanger sequencing (**Fig. 3D-E**). Further validation was carried out in PBMCs from 4 patients with atopic dermatitis and serum IgE levels between 2370 and 6350 kIU/L. *Bona fide* IgE^+^ MBC, confirmed with Sanger sequencing, were identified in 2 of these 4 patients at a frequency of 0.0015% from total B cells (**Fig. 3F-G**).

### IgE^+^ MBCs are undetectable in PN-allergic patients

With our enhanced detection method, we conducted analysis on PBMCs of 20 donors that included PN-allergic (n=9; mean serum total IgE of 196 [11-890] kIU/L) and non-allergic patients (n=10) (**Table S1**). We detected similar frequencies of putative IgE^+^ MBCs (% from total B cells = 0.0019 PN-allergic, 0.0046 non-allergic). However, in all instances there was no IgE amplification (**Table 1**). To ensure that IgE^+^ MBCs were not being undetected through our exclusion of CD27^−^ cells, we sorted CD27^−^ IgE^+^ MBCs as it has been speculated that MBCs arising extra-follicularly (*14*) may not gain the canonical MBC marker, CD27 (*35*). Similarly, no PCR amplification occurred with IgE primers (data not shown). Furthermore, we investigated the possibility that the polyclonal anti-IgG antibody could bind non-specifically to IgE^+^ MBCs due to serum IgG or IgA bound to MBCs, thus masking IgE^+^ MBCs cells in the IgG^+^ or IgA^+^ populations. Here, we stained for IgA and IgG on the same fluorophore and flow-sorted class-switched MBCs that were positive for IgE (**Fig. S1**). The frequency of this population was 0.074% and the genetic analysis demonstrated that these MBCs were of a non-IgE identity that presumably bound secreted IgE.

**Supplementary Figure 1.**
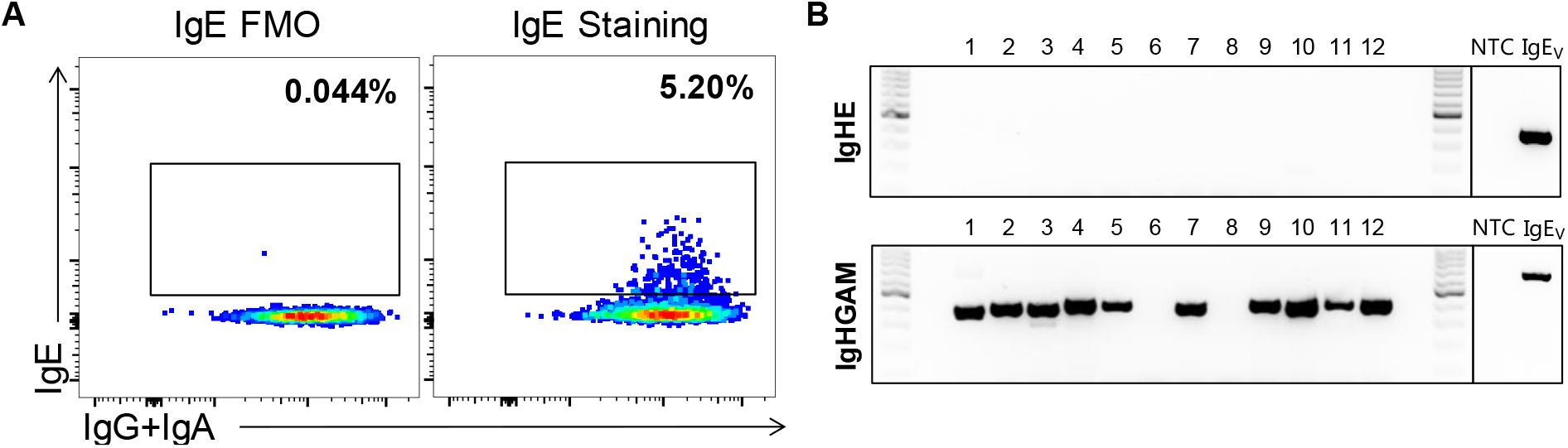
Assessment of cytotropic staining of IgG and/or IgA in IgE^+^ MBCs. **(A)** Cytometry detection and sorting of class-switched memory B cells that stained positive for IgE and IgG/IgA; **(B)** BCR amplification with primers specific for IgE (IgHE) or a mix specific of IgG, IgA and IgM (IgHGAM) of single-sorted cells. Data are representative of 2 independent experiments (2 donors per experiment and 18 cells single-sorted per donor).

**Supplementary Table 1.**
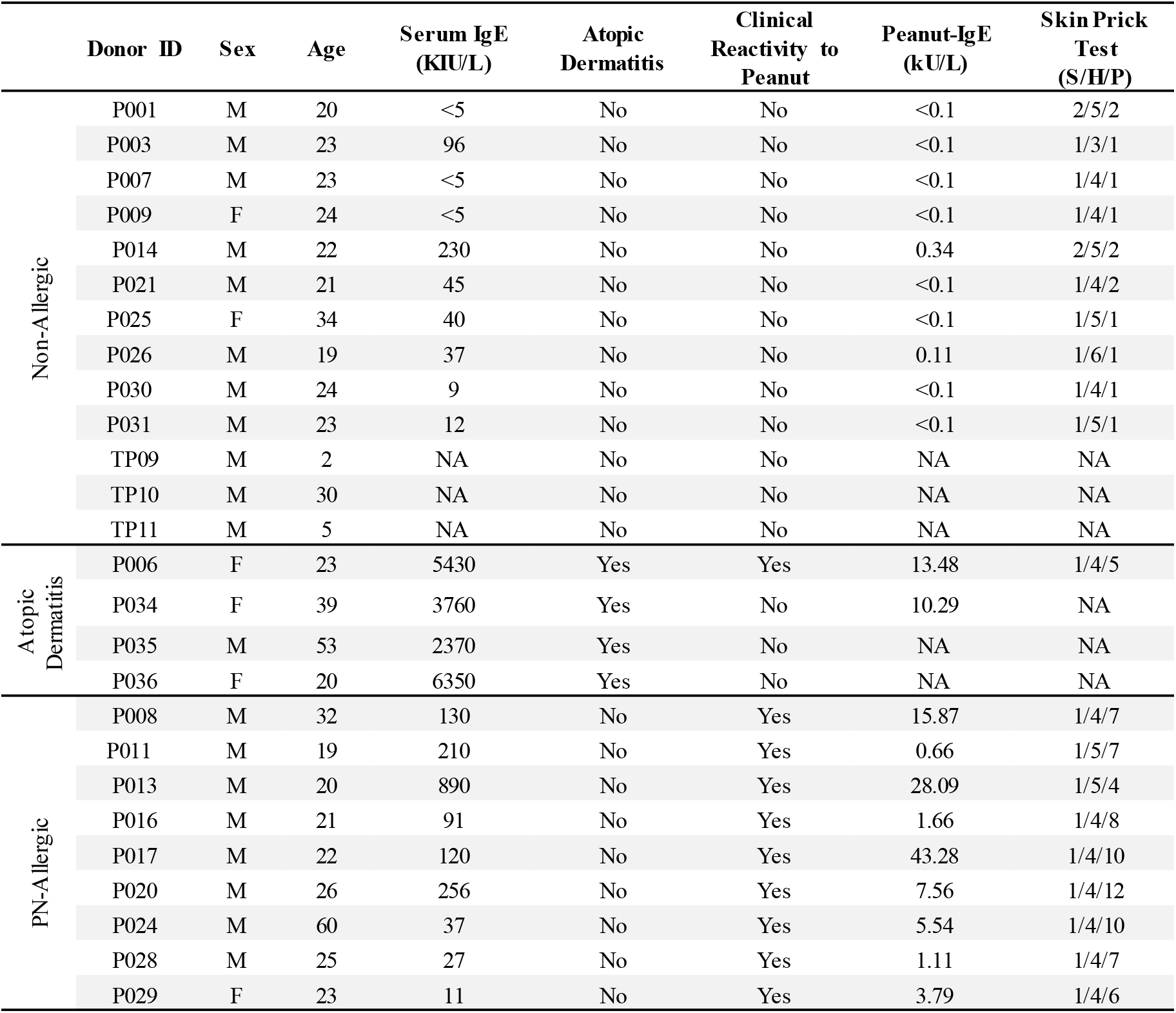
Patient’s profiles

**Table 1.** Quantification of IgE^+^ MBCs in healthy and allergic donors

| Tissue | Donor ID | Allergic Status | MNCs | Purified B Cells | Events in CD20<sup>+</sup> CD38<sup>lo-med</sup> Gate | Events in IgE Gate | Sorted Cells | IGHE Amplification |
| --- | --- | --- | --- | --- | --- | --- | --- | --- |
| Blood | P001 | - | 250,000,000 | 9,540,000 | 600,304 | 21 | 6 | 0 |
|  | P003 | - | 123,000,000 | 8,640,000 | 169,267 | 29 | 12 | 0 |
|  | P007 | - | 125,000,000 | 2,685,000 | 645,055 | 5 | 3 | 0 |
|  | P009 | - | 98,600,000 | 4,140,000 | 325,734 | 12 | 12 | 0 |
|  | P014 | - | 109,000,000 | 2,820,000 | 605,535 | 8 | 8 | 0 |
|  | P021 | - | 173,000,000 | 26,000,000 | 440,963 | 2 | 1 | 0 |
|  | P025 | - | 86,600,000 | 1,401,000 | 246,090 | 20 | 20 | 0 |
|  | P026 | - | 94,200,000 | 1,494,000 | 250,018 | 14 | 10 | 0 |
|  | P030 | - | 78,800,000 | 1,128,000 | 132,157 | 4 | 4 | 0 |
|  | P031 | - | 85,400,000 | 945,000 | 172,260 | 4 | 4 | 0 |
|  | P008 | PN | 125,000,000 | 8,160,000 | 335,641 | 20 | 20 | 0 |
|  | P011 | PN | 250,000,000 | 7,150,000 | 520,021 | 6 | 3 | 0 |
|  | P013 | PN | 210,000,000 | 5,450,000 | 690,015 | 13 | 12 | 0 |
|  | P016 | PN | 125,000,000 | 1,068,000 | 143,759 | 5 | 5 | 0 |
|  | P017 | PN | 125,000,000 | 1,467,000 | 213,345 | 11 | 10 | 0 |
|  | P020 | PN | 124,000,000 | 3,900,000 | 537,856 | 4 | 1 | 0 |
|  | P024 | PN | 71,800,000 | 1,026,000 | 166,313 | 3 | 1 | 0 |
|  | P028 | PN | 94,800,000 | 1,440,000 | 157,340 | 1 | 1 | 0 |
|  | P029 | PN | 54,000,000 | 1,467,000 | 277,359 | 2 | 2 | 0 |
| Tonsils | TP-9 | - | 10,000,000 | - | 204,125 | 10 | 7 | 0 |
|  | TP-10 | - | 10,000,000 | - | 68,461 | 15 | 12 | 0 |
|  | TP-11 | - | 10,000,000 | - | 56,127 | 6 | 6 | 0 |

## Discussion

Persistent memory underlies the lifelong maintenance of some food allergies such as those to PN, tree nuts, fish and shellfish. As clinical reactivity is largely mediated by IgE, vast efforts have been dedicated to identifying IgE^+^ MBCs and their role in this process. In murine systems, flow cytometry, immunofluorescence, genetic analysis, *in vitro* cultures, IgE reporter mice and functional models of adoptive transfer (*17, 18, 21, 36–38*) have been used to study IgE^+^ MBCs. In humans, putative IgE^+^ MBCs have been studied mainly by flow cytometry (*24, 27, 28*), occasionally validated via genetic analysis in bulk (*14*) to roughly infer the frequency of the different BCR-transcripts expression in B cell populations (*39*). However, increasing evidence in mice and humans has questioned the relevance, even the existence, of IgE^+^ MBCs as the true reservoir of IgE-secreting cells (*8, 15, 19, 20, 31, 40*). Here, we set out to ascertain the true nature of putative IgE^+^ MBCs identified by flow cytometry through genetic validation at the single cell level.

Our enhanced step-wise exclusion protocol eliminated spurious events in the IgE gate, which coincided with the disappearance of the double-negative population when staining IgG, IgA and IgE BCRs of class-switched MBCs. The fidelity of this protocol was validated in studies using PBMCs cultures stimulated with IL-4 and anti-CD40, which induces IgE class switching. Molecular validation at the single cell level demonstrated *IgE* transcripts and, thus, confirmed the identification those cells as *bona fide* IgE^+^ MBCs. PBMCs from 4 patients with atopic dermatitis were investigated. *Bona fide* IgE^+^ MBCs were identified in 2 of these patients. While their frequency was extremely rare, 0.0015% of B cells, their detection further validates the methodologies used. We then extended this analysis to PBMC samples from 19 individuals, including 9 PN-allergic patients and 10 healthy, non-allergic controls as well as tonsils from 3 non-allergic subjects. Using our enhanced flow cytometry protocol and genetic validation approach, we did not detect IgE^+^ MBCs in any of the samples from non-allergic controls or patients with PN allergy, only a residual population at a frequency comparable to that of the FMO.

The value of scientific knowledge relies to a large extent on the fidelity of the tools used to generate such knowledge. In this context, we provide a validated method to identify *bona fide* IgE^+^ MBCs. Our data demonstrate the extreme rarity of these cells in the circulation of allergic patients-at least orders of magnitude lower than previously reported (*14, 24*)- and are in agreement with human genetic studies that reported few IgE transcripts in circulation but without unambiguously defining the B cell phenotype (MBC, plasmablast, *etc*.) (*41, 42*). This finding strengthens the concept that the reservoir of IgE-secreting cells resides in MBCs of a non-IgE isotype and, as such, informs future research directions. It is possible that tissues from allergic subjects could harbour IgE^+^ MBCs; this remains both a challenge and a venue for future efforts, Nevertheless, it is evident that the proposal that circulating IgE^+^ MBCs could be a clinical marker for allergic disease is unproven.

## Materials and Methods

### Flow cytometry

Antibodies were obtained from BioLegend (San Diego, California), Columbia Biosciences (Frederick, Maryland), BD Biosciences (San Jose, California), Miltenyi Biotec (Bergisch Gladbach, Cologne), eBioscience (Carlsbad, California) or ThermoFisher Scientific (Waltham, Massachusetts): CD38-phycoerythrin (PE)-Cy7 (clone HIT2); IgE-allophycocyanin (APC) (Columbia Biosciences SKU: D3-110-E); IgG-PE (clones G18-145 and HP6017, and ThermoFisher Scientific Catalog #12-4998-82); IgA-PE (clone IS11-8E10); IgA-APC (clone IS11-8E10); IgM-Brilliant Violet (BV) 510 (clone MHM-88); IgD-BV421 (clone IA6-2); IgG-biotin (ThermoFisher Scientific Catalog #A18815); CD20-Alexa Fluor700 (clone 2H7); CD27-FITC (clone O323). In all assays 1 x 10^6^ cells were first incubated with Human TruStain FcX (Fc Receptor Blocking Solution, Biolegend) or anti-human CD32 (FcγRII Blocker, StemCell Technologies) for 15 minutes on ice to block non-specific staining and then incubated with fluorochrome-conjugated antibodies for 30 minutes on ice and protected from light. When IgG-biotin was used to label IgG^+^ cells, cells were incubated for an additional 30 minutes with streptavidin-PE (BioLegend) on ice and protected from light. Dead cells were excluded using the fixable viability dye eFluor780 (eBioscience) and by gating on singlets. FMO were used for gating. Data were acquired on a Fortessa (BD Biosciences) and analyzed with FlowJo software (TreeStar, Ashland, Ore), and single cells were sorted on a MoFlo XDP Cell Sorter (Beckman Coulter).

### PCR amplification

Single cells were sorted into 96-well PCR plates (Thermofisher) containing 20 units RNasin^®^ Ribonuclease Inhibitors (Promega) 2 μL First Strand buffer (Thermofisher), and nuclease-free water to a volume of 10 μL per well. The cells are centrifuged at 4°C at 550 *g* for 1 minute and immediate placed in −80°C freezer. Next, heat lysis was performed by adding 3 μL Nonidet^®^ P-40 Substitute (G-Biosciences) and 150 ng of random hexamers (Thermofisher). The reaction was performed at 65°C for 10 minutes and then 25°C for 3 minutes. All thermocycler reactions were done using Mastercycler^®^ pro S (Eppendorf). cDNA was synthesized as previously described. Briefly, 2 μL 5X First Strand buffer (Thermofisher), 2 μL of 0.1 M Dithiothreitol (Qiagen), 1 μL of 10 mM each dNTP, and 0.5 μL SuperScript III (Thermofisher) was added to the plate containing the heat lysis contents (final volume 19.5 μL). Reverse transcription was performed at 37°C for 1 hour and then 70°C for 10 minutes.

IgH amplification was accomplished using a two-step nested PCR as previously described. Briefly, a mix of 6 forward primers and either a reverse primer specific to *IGHE* or a pool of reverse primers specific to *IGHM, IGHA* and *IGHG* were used (*IGHM, IGHA, IGHG* primer sequences found in Tiller *et al*.(*29*), IGHE primer sequences found in Berkowska *et al*. (*14*). The first PCR reaction contained 8 μL of cDNA mixture, 1 unit of HotStar Plus Taq (Qiagen), 200 nM of each primer, 400 μM dNTP (Thermofisher), 5 μL 10X PCR buffer (Qiagen), and nuclease-free water to a final volume of 50 μL. The reaction was performed starting with 3 cycles of pre-amplification of 95°C for 45 seconds, 45°C for 45 seconds, 72°C for 45 seconds, followed by 30 cycles of 94°C for 45 seconds, 50°C for 45 seconds, 72°C for 1 minute and 45 seconds, followed by a final extension of 72°C for 10 minutes. PCR 2 was performed using a new nested pool of forward primers as well as nested reverse primers specific to *IGHE* or a combination of *IGHM, IGHG, IGHA* (*IGHM, IGHG, IGHA* primer sequences found in Tiller *et al*. (*29*), IGHE primer sequences found in Berkowska *et al*.(*14*). The PCR reaction contained 4 μL of PCR 1 product, 5 μL of 10X Pfu buffer (Agilent) 1.25 μL of 10 mM dNTP (Thermofisher), 400 nM of each primer, 1.25 units Pfu polymerase (Agilent), and nuclease-free water to a final concentration of 50 μL. The reaction was performed for 30 cycles at 94°C for 45 seconds, 50°C for 45 seconds, 72°C for 1 minute and 45 seconds, followed by a final extension of 72°C for 10 minutes.

The second PCR product was visualized on a 1.5% agarose gel and the expected band size was approximately 400 bp. Amplified IgH sequences were enzymatically purified using ExoSAP-IT™ PCR Product Cleanup Reagent (Thermofisher) and subsequently Sanger sequenced using the forward and reverse primers (GENEWIZ). The sequences were analyzed using IMGT/HighV-QUEST (http://imgt.org/HighV-QUEST) for V, D, and J sequences with the highest identity, as well as nucleotide and amino acid mutations from their germline sequences.

### Generation of a DNA vector containing a human IgE backbone (IgEV)

To generate a human IgE, we started with a heavy chain IgG1 vector (gifted by Michel C. Nussenzweig) previously modified to include an Ara h 2 variable chain (*33*). The human ε constant region was amplified from an anti-OVA human IgE vector (*43*) with primers (5’ TTTT GTCGAC GGCGCACCA 3’ and 5’ TTTT AAGCTT CTCAATGGTGGTGATGTTTA 3’) to add flanking SalI and HindIII restriction enzyme sites. The human ε constant region then replaced the γ1 constant region under the CMV provider using the SalI and HindIII restriction enzyme sites. Sanger sequencing confirmed successful insertion of the ε constant region.

### Study population

A cohort of 10 PN-allergic and 10 non-allergic blood donors were recruited from McMaster University (Hamilton, ON). Allergy to PN was ascertained by PN-specific IgE ImmunoCap^®^ performed at LRC Hamilton (McMaster Children’s Hospital), and by skin prick test. PN-allergic individuals were considered for inclusion with PN-specific serum IgE levels >0.35 kU/L and skin prick test ≥3 mm greater than saline control. Total serum IgE was quantified by IMMAGE^®^ 800 (Beckman Coulter) performed at LRC Hamilton for a general measure of atopy. We recruited an additional 4 participants with total serum IgE levels > 2300 kIU/L and received 3 tonsil discards from individuals undergoing routine tonsillectomies. Exclusion criteria for all recruited donors included: allergen immunotherapy, previous or current omalizumab (Xolair^®^) treatment, other systemic immunomodulatory treatments (*i.e*., rituximab), or autoimmune/immunodeficiency diseases. Patient demographics and allergic indicators are summarized in **Table S1**. All donors were recruited with written consent and ethical approval from Hamilton Integrated Research Ethics Board (HiREB).

### Mononuclear cell isolation and B cell enrichment

Up to 80 mL of peripheral blood was collected into heparinized tubes (BD) and tonsils were crushed into a single-cell suspension. PBMCs were isolated via Ficoll-Paque (GE Healthcare) density gradient centrifugation. Immediately following, B cells were isolated from PBMCs using a negative selection magnetic separation kit (19054, StemCell Technologies) with at least 70% purity.

### PBMC culture

PBMCs were cultured in RPMI 1640 (Gibco) supplemented with 10% human AB serum (Corning), 10 mM HEPES, 0.1 mM non-essential amino acids, 1 mM sodium pyruvate, 55 μM 2-mercaptoethanol, 1% L-glutamine, and 1% penicillin-streptomycin. Cells were plated at a density of 1.5 x 10^6^ per mL in 24-well plates. Stimulated cells were treated with 68.7 ng/mL (8000IU) IL-4 (Sigma-Aldrich) and 5 μg/mL anti-CD40 (BioXCell) on day 1. Cells were incubated at 37°C and 5% CO_2_ for the duration of culture. On days 4 and 8 of culture, 1 mL of cell-free supernatant was withdrawn and replaced with fresh media. Supernatant was also withdrawn on day 11 of culture and stored at −80°C for later analysis of total IgE by ELISA. After 11 days in culture, cells were harvested and IgE-secreting cells were quantified using ELISPOT.

### ELISA and ELISPOT

For total IgE ELISA, MaxiSorb plates (ThermoFisher Scientific) were coated with 0.5 μg/ml antihuman IgE (555894, BD Pharmingen) in carbonate-bicarbonate buffer overnight at 4°C. Coated wells were blocked with 5% skim milk in PBS for 2 hours at room temperature, followed by 3 washes (1x PBS and 0.05% Tween 20). Cell-free supernatant samples and a serial dilution of purified human IgE (401152, Calbiochem) were incubated overnight at 4°C. Wells were washed 3 times and incubated with 1 μg/ml biotinylated anti-human IgE (A18803, Invitrogen) in 1% skim milk for 2 hours at room temperature. Subsequently, wells were washed 3 times and incubated with alkaline-phosphatase streptavidin for 1 hour at room temperature. Plates were developed with p-nitrophenyl phosphate and the reaction was stopped with 2N NaOH. Optical density was measured at 405 nM (Multiskan^−^FC, Thermo Scientific).

A commercially available ELISPOT kit (3810-2H, Mabtech) was used for the detection of IgE-secreting cells. On day 11 of culture, samples were plated in duplicate at 4 x 10^6^ cells/mL. Plates were imaged with ImmunoSpot^®^ S6 Analyzer and spots were counted independently by 2 blinded investigators.

## Acknowledgments

We acknowledge all the members from the laboratories of Drs. Jordana and Shreffler for scientific input and technical help. We thank Joshua F.E. Koenig for critical review of the manuscript.

## Author contributions

R.J-S. designed the study. R.J-S., Y.E., K.B., P.S. planned and performed the experiments, analyzed the data, and wrote the manuscript. S.W., D.D.S. and H.L. participated in patient recruitment, clinical data, and biological sample collection. S.U.P, W.G.S, S.W. and M.J. raised funding, provided feedback and reviewed the manuscript. M.J. supervised the project.

## Data and materials availability

If data are in an archive, include the accession number or a placeholder for it. Here also include any materials that must be obtained through an MTA. Acknowledgments follow the references and notes but are not numbered.

